# Aquaporin 4 is differentially increased and depolarized in association with tau and amyloid-beta

**DOI:** 10.1101/2022.04.26.489273

**Authors:** Vasil Kecheliev, Leo Boss, Upasana Maheshwari, Uwe Konietzko, Annika Keller, Daniel Razansky, Roger M. Nitsch, Jan Klohs, Ruiqing Ni

**Affiliations:** Institute for Regenerative Medicine, University of Zurich, Zurich, Switzerland; Department of Neurosurgery, Clinical Neuroscience Center, Zürich University Hospital, Zurich, Switzerland; Zentrum für Neurowissenschaften Zurich, Zurich, Switzerland; Institute for Biomedical Engineering, ETH Zurich & University of Zurich, Zurich, Switzerland

**Keywords:** Alzheimer’s disease, amyloid-beta, aquaporin 4, astrocyte, magnetic resonance imaging, perfusion, tau

## Abstract

Neurovascular-glymphatic dysfunction plays an important role in Alzheimer’s disease and has been analyzed mainly in association with amyloid-beta (Aβ) pathology. The neurovascular-glymphatic link with tauopathies has not been well elucidated. Here, we aimed to investigate the alterations in the neurovasculature and map the aquaporin 4 (AQP4) distribution and depolarization associated with tau and Aβ. Perfusion, susceptibility weighted imaging and structural magnetic resonance imaging (MRI) were performed in the pR5 P301L mouse model of 4-repeat tau and the arcAβ mouse model of amyloidosis. Immunofluorescence staining was performed using antibodies against AQP4, CD31, astroglia (GFAP, s100β), phospho-tau (AT-8) and Aβ (6E10) in brain tissue slices from P301L, arcAβ and nontransgenic mice. P301L mice showed regional atrophy, preserved cerebral blood flow and reduced cerebral vessel density compared to nontransgenic mice, while arcAβ mice showed cerebral microbleeds and reduced cerebral vessel density. AQP4 depolarization and peri-tau enrichment in the hippocampus and increased AQP4 levels in the forebrain and hippocampus were detected in P301L mice compared to nontransgenic mice. In comparison, cortical AQP4 depolarization and cortical/hippocampal peri-plaque increases were observed in arcAβ mice. Increased s100β-GFAP fluorescence intensities indicative of reactive astrocytes were detected surrounding tau inclusions in P301L mice and Aβ plaques in arcAβ mice. In conclusion, we observed a divergent region-specific AQP4 increase and association with phospho-tau and Aβ pathologies.

## Introduction

Astrocytes are essential for maintaining homeostasis, synaptic plasticity and the inflammatory response in the central nervous system [1] and show disease-associated profiles in AD [2–5]. Similar to previous studies in animal studies, glymphatic drainage has been demonstrated by MR imaging in live human brains [6, 7]. Contribution of astrocytic tau pathology to synapse loss in primary tauopathies, including progressive supranuclear palsy and corticobasal degeneration [8]. The glymphatic system plays an important role in neurodegenerative diseases, including Alzheimer’s disease (AD) and primary tauopathy [9–15]. Dysfunction of the glymphatic system reduces the removal of amyloid-beta (Aβ) and tau proteins [16–18]. Aquaporin-4 (AQP4) constitutes a critical part of the glymphatic system. In the central nervous system, AQP4 is located in either the perivascular endfoot or neuron-facing membrane domains of astrocytes [19, 20]. AQP4 facilitates the movement of cerebrospinal fluid (CSF) and its mixing with interstitial fluid and is essential in the clearance of accumulated toxic Aβ and tau aggregates from the brain [21–23]. Loss of perivascular AQP4 localization has been shown to be associated with cognition and Aβ plaque and cerebral amyloid angiopathy (CAA) in patients with AD [24, 25]. *AQP4* polymorphisms predict Aβ burden, cognition and clinical progression in the AD spectrum [26, 27] and moderate the relationship between sleep and brain Aβ burden [28]. Thus, AQP4 subcellular relocalization in water homeostasis of the central nervous system has been a potential drug target for neurodegenerative diseases [29–34]. Emerging evidence suggests an association of AQP4 with tau, such as a marker of the ageing-related tau astroglial response [35]. Transcriptional network analysis of human astrocytic endfoot genes has revealed region-specific associations with dementia status and tau pathology [36]. A higher CSF level of AQP4 was observed in presymptomatic primary tauopathy frontotemporal dementia (FTLD) mutation carriers than in healthy controls [37].

Impaired perfusion, capillary function and neurovascular dysfunction play an important role in patients with AD and primary tauopathy as well as in animal models [38–44]. Cerebral blood flow (CBF) has been shown to be a driver of glymphatic function [45, 46]. Restriction of regional CBF reduces arterial wall pulsatility, thus reducing CSF-interstitial fluid exchange in the brain. In turn, inhibition of AQP4 significantly increased rCBF in wild-type mice [47]. Increased cerebral vascularization and decreased water exchange were detected across the blood–brain barrier (BBB) in AQP4 knockout mice [48]. A few recent studies have used noninvasive magnetic resonance imaging (MRI) of brain clearance pathways using multiple echo time arterial spin labelling (ASL) [49–53].

Many studies have demonstrated the association of Aβ and AQP4, but the spatial association between tau and AQP4 is not fully clear. Here, we evaluated the alterations in cerebrovascular function using high-field MRI and atroglial AQP4 distribution in relation to tau and Aβ by immunofluorescence staining in 4-repeat tau (pR5 p301L) and amyloidosis (arcAβ) mouse models.

## Materials and methods

### Animal models

Eleven transgenic arcAβ mice overexpressing the human APP695 transgene containing the Swedish (K670N/M671L) and Arctic (E693G) mutations under the control of the prion protein promoter and 11 age-matched nontransgenic littermates (NTL) of both sexes were used in this study (16-20 months of age). The arcAβ mouse model of cerebral amyloidosis, which exhibits strong CAA [54–57], displays decreased microvessel density [58], blood brain barrier leakage [54], vascular remodelling [59], and cerebral microbleeds (CMBs) [60]. Transgenic B6.Dg-Tg(Thy1.2-TauP301L) mice have been engineered to express the human 4 repeat tau isoform under the control of the murine Thy1.2 promoter. This pR5 tau mouse model has shown alterations in tau accumulation and white matter integrity in previous studies [61–65]. After generation, mice were backcrossed with C57BL/6J mice for > 20 generations and maintained on a C57BL/6J background. Neuroinflammatory changes and atrophy were reported in the cervical spinal cord of P301L mice [66]. A total of 8 hemizygous P301L mice and 10 NTLs of 10-12 months of age were used. Animals were housed in ventilated cages inside a temperature-controlled room under specific pathogen-free conditions and under a 12-hour dark/light cycle. Pelleted food (3437PXL15, CARGILL) and water were provided *ad libitum*. Paper tissue and red Tecniplast^®^ mouse house (Tecniplast, Italy) shelters were placed in cages for environmental enrichment. All experiments were performed in accordance with the Swiss Federal Act on Animal Protection and were approved by the Cantonal Veterinary Office Zurich (permit number: ZH082/18, ZH162/20).

### Perfusion MRI in P301L mice

Perfusion MRI was performed using an arterial spin labelling technique as described previously [62, 67]. Perfusion MRI scans were performed on a 7/16 small animal MR Pharmascan (Bruker Biospin GmbH, Ettlingen, Germany) equipped with an actively shielded gradient set of 760 mT/m with an 80 μs rise time and operated by a Paravision 6.0 software platform (Bruker Biospin GmbH, Ettlingen, Germany). Mice were anesthetized with an initial dose of 4% isoflurane (Abbott, Cham, Switzerland) in an oxygen/air (200/800 μl/min) mixture, and anesthesia was maintained at 1.5 % isoflurane in an oxygen/air (100/400 μl/min) mixture. Mice were placed in the prone position on a water-heated support to keep body temperature within 36.5 ± 0.5 °C. Body temperature was monitored with a rectal temperature probe. rCBF maps from pulsed arterial spin labelling data were computed as previously described. Regions of interest (ROIs) were evaluated in the cerebral cortex and hippocampus based on T_2_-weighted anatomical scans. rCBF in ROIs was calculated from both T_1_ values using MATLAB R2019b (Mathworks, United States) [62, 68]. The Allen mouse brain atlas was used as an anatomical reference for scan positioning and data analysis [69].

### *In vivo* susceptibility weighted imaging (SWI) MRI in arcAβ mice

MRI was carried out on a Bruker Biospec 94/30 (Bruker Biospin GmbH, Ettlingen, Germany) small animal MR system at 400 MHz equipped with a BGA 12AS HP gradient system with a maximum gradient strength of 400 mT/m and minimum rise time of 70 μs [70] operated under Paravision 6. The system was equipped with a cryogenic 2 × 2 phased-array cryogenic mouse head surface coil (Bruker BioSpin AG, Fällanden, Switzerland). Mice were anesthetized with an initial dose of 4 % isoflurane in an oxygen/air (200:800 ml/min) mixture and were maintained at 1.5 % isoflurane. Mice were endotracheally intubated and mechanically ventilated with 90 breaths per minute (bpm), applying a respiration cycle of 25 % inhalation and 75 % exhalation (MRI-1 Volume Ventilator CWI Inc., Ardmmore, PA, USA). The tail vein was cannulated with a 30-gauge needle for subsequent intravenous (i.v.) injection (0.3 mm × 13 mm; BD Microlance, Drogheda, Ireland). Body temperatures were monitored with a rectal temperature probe (MLT 415, AD Instruments, Spechbach, Germany) and kept at 36.5 ± 0.5 °C on a water-heated holder (Bruker Biospin AG, Switzerland). Anatomical reference data were acquired in coronal and sagittal directions. After standard adjustments, gradient echo images in 3 orthogonal directions were acquired at Bregma 1.52 mm. Data for SWI were acquired using a flow-compensated gradient echo sequence. The image parameters were as follows: axial, how many slices, slice gaps, time to echo (TE) = 5 ms; repetition time (TR) = 630 ms; flip angle (FA) = 16°; field-of-view (FOV) = 4 cm; matrix size = 512 × 512; spatial resolution = 0.078×0.078×0.34 mm/pixel; and total scan time = 8 minutes. SW and phase images were computed using the SWI processing module in ParaVision 6.0.1 (Bruker, Ettlingen, Germany) with Gauss broadening □=□ 1 mm and mask weighting □=□4. All SW images were compared with their phase image counterparts to ensure that the signals were paramagnetic (presence of CMBs). CMBs were counted manually twice for isolated circular shapes in all slices using MRIcron (University of South Carolina, USA) as described previously [71].

### *Ex vivo* structural MRI in P301L mice

After *in vivo* imaging, P301L and NTL were intracardially perfused under deep anesthesia (ketamine/xylazine/acepromazine maleate (75/10/2 mg/kg body weight, i.p. bolus injection) with 0.1 M phosphate buffered saline (PBS, pH 7.4). The heads were postfixed in 4 % paraformaldehyde in 0.1 M PBS (pH 7.4) for 6 days and stored in 0.1 M PBS (pH 7.4) at 4 °C. Brains were not removed from the skull, which has been shown previously to preserve cortical and central brain structure. The heads were placed in a 15 ml centrifuge tube filled with perfluoropolyether (Fomblin Y, LVAC 16/6, average molecular weight 2700, Sigma–Aldrich, U.S.A.). Data were acquired on a BioSpec 94/30 with a cryogenic 2×2 radio frequency phased-array surface coil (overall coil size 20×27 mm^2^, Bruker BioSpin AG, Fällanden, Switzerland) with a coil system operating at 30□K (Bruker BioSpin AG, Fällanden, Switzerland) for reception used in combination with a circularly polarized 86□mm volume resonator for transmission.

### Immunofluorescence staining

For immunofluorescence staining, mice were perfused under ketamine/xylazine/acepromazine maleate anesthesia (75/10/2 mg/kg body weight, i.p. bolus injection) with ice-cold 0.1 M PBS (pH 7.4) and in 4 % paraformaldehyde in 0.1 M PBS (pH 7.4), fixed for 24 h in 4 % paraformaldehyde (pH 7.4) and then stored in 0.1 M PBS (pH 7.4) at 4 °C. Coronal brain sections (40 μm) were cut around bregma −2-0 mm and stained with anti-Aβ antibody 6E10 anti-phospho-tau antibodies AT-8, AT-100, antibodies against AQP4, CD31 for platelet endothelial cell adhesion molecule 1 (PECAM1), glial fibrillary acidic protein (GFAP), S100 calcium binding protein β (s100β), cluster of differentiation 68 (CD68), and ionized calcium-binding adapter molecule 1 (Iba1) as described previously [62, 67, 70]. Sections were counterstained using 4’,6-diamidino-2-phenylindole (DAPI) and mounted with VECTASHIELD antifade fluorescent mounting media (details in **STable 1**). Hematoxylin and eosin (H&E) staining was performed on arcAβ mouse brain slides. The brain sections were imaged at 20× magnification using an Axio Oberver Z1 (Zeiss, Germany) with the same acquisition settings for all brain slices and a Leica SP8 confocal microscope (Leica, Germany). The images were analyzed blindly using Qupath [72] and FIJI (NIH, USA). For Iba1-6E10 staining, the images were examined using Leica DM 4000B microscopy with Olympus DP71 digital camera and VIS (Visiopharm Integrator System) version 4.4.6.9 software at 5 ×, 10 ×, 20 × and 40 × magnification.

Perivascular polarization of AQP4 was measured as previously described [52, 73]. Briefly, the median immunofluorescence intensity of perivascular (CD31) regions was measured. A threshold analysis was then used to measure the percentage of the region exhibiting AQP4 immunofluorescence greater than or equal to perivascular AQP4 immunofluorescence (AQP4 cover area %). For measurement of AQP4 association with tau and amyloid, the background corrected mean fluorescence immunoreactivity of AQP4 labelling was determined within manually drawn ROIs in the peri-tau or peri-plaque regions and divided by the mean intensity within nonplaque parenchymal areas as previously described [74].

### Statistics

Group comparisons in multiple brain regions were performed by using two-way ANOVA with Bonferroni’s *post-hoc* analysis (GraphPad Prism 9). Comparisons between the number of CMBs of arcAβ and NTL and the cortical thickness of P301L and NTL were performed by using a Mann–Whitney test. All data are presented as the mean ± standard deviation. Significance was set at \**p* < 0.05.

## Results

### Preserved rCBF, reduced cortical thickness and reduced cortical and hippocampal vessel density in P301L mice

First, we examined the functional and structural changes in the 10-month-old P301L mouse model compared to NTL. Perfusion ASL-MRI showed comparable rCBF in the cortex (104.5 ± 16.4 vs 104.3 ± 24.4) and hippocampus (116.5 ± 14.5 vs 105.5 ± 16.4) of P301L mice (n = 4) compared to NTL (n = 4) (**Figs. 1a, e**). Regional brain atrophy indicated by reduced cortical thickness (0.7 ± 0.1 vs 1.0 ± 0.2, p = 0.031) was detected in P301L mice (n = 4) compared to NTL mice (n = 8) (**Figs. 1b. f**). The degree of atrophy observed in the pR5 P301L mice was much milder than that in rTg4510 mice from previous studies [75, 76]. Immunohistochemical staining using CD31 showed reduced vessel density in the forebrain (2.4 ± 0.2 vs 4.3 ± 0.9, p = 0.0039) and CA1 of the hippocampus (2.4 ± 0.3 vs 4.3 ± 0.4, p = 0.0048) of P301L mice (n = 3) compared to NTL (n = 3) (**Figs. 1c, d, g**). The accumulation of phospho-tau inclusions (AT-8 positive) in the forebrain and hippocampus of P301L mice was detected. No specific detection of AT-8 or AT-100 immunoreactivity was detected in the brains of NTL. Hippocampal atrophy was observed in the brain tissue slices of P301L mice (**SFig. 1**).

**Fig. 1.**
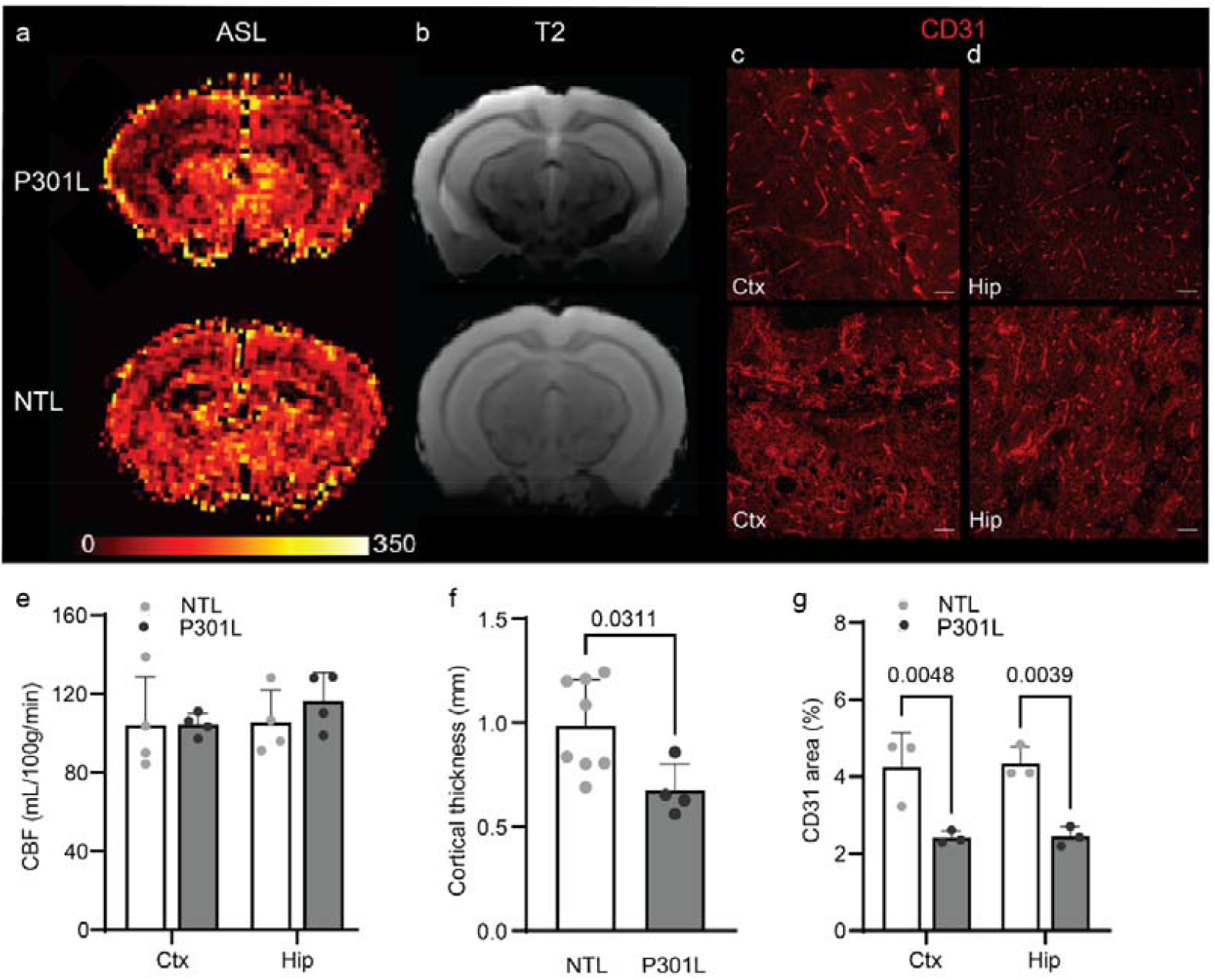
Preserved cerebral perfusion, regional atrophy and reduced vessel density in the P301 L mouse brain. (**a, e**) Representative coronal regional cerebral blood flow (rCBF) map of 12-month-old P301L (upper row, n = 4) and nontransgenic littermate (NTL) mice (lower row, n = 4); CBF scale: 0-350 ml/100 g/min. (**b, f**) T_2_ MRI at 9.4 T showing reduced cortical thickness of P301L compared to NTL mice; (**c, d, g**) CD31 staining showing reduced vessel density on the cortex and hippocampus of P301L compared to NTL mice. Scale bar = 20 μm. Data are presented as the mean ± standard deviation. ASL, arterial spin labelling.

### AQP4 increase and depolarization and peri-tau enrichment in P301L mice

Next, we performed triple staining using AQP4/CD31/AT8 (phospho-tau) and AQP4/GFAP/AT8 to investigate the association and status of AQP4 with pathology in P301L mice. AQP4 staining immunoreactivity was homogenous in the brains of NTL mice (**Fig. 2a, b**, **SFig. 2**). Colocalization of tau inclusions with CD31-labelled endothelial cells and with AQP4 was observed in the forebrain and hippocampus of P301L mice (**Figs. 2c-f**). The level of AQP4 immunofluorescence (absolute value) was significantly higher in the forebrain (39.8 ± 2.1 vs 22.8 ± 1.6, approx. 75 % increase, p < 0.0001) and hippocampal regions (30.0 ± 2.6 vs 21.6 ± 1.0, approx. 40 % increase, p = 0.0013) of P301L (n = 3) than in NTL mice (n = 3) (**Figs. 2g**). The peri-tau level of AQP4 was increased compared to that of the parenchymal (peri-tau/parenchymal ratio approx. 1.3 ± 0.05) in the hippocampus of P301L mice but not in the forebrain (**Fig. 2h**).

**Fig. 2.**
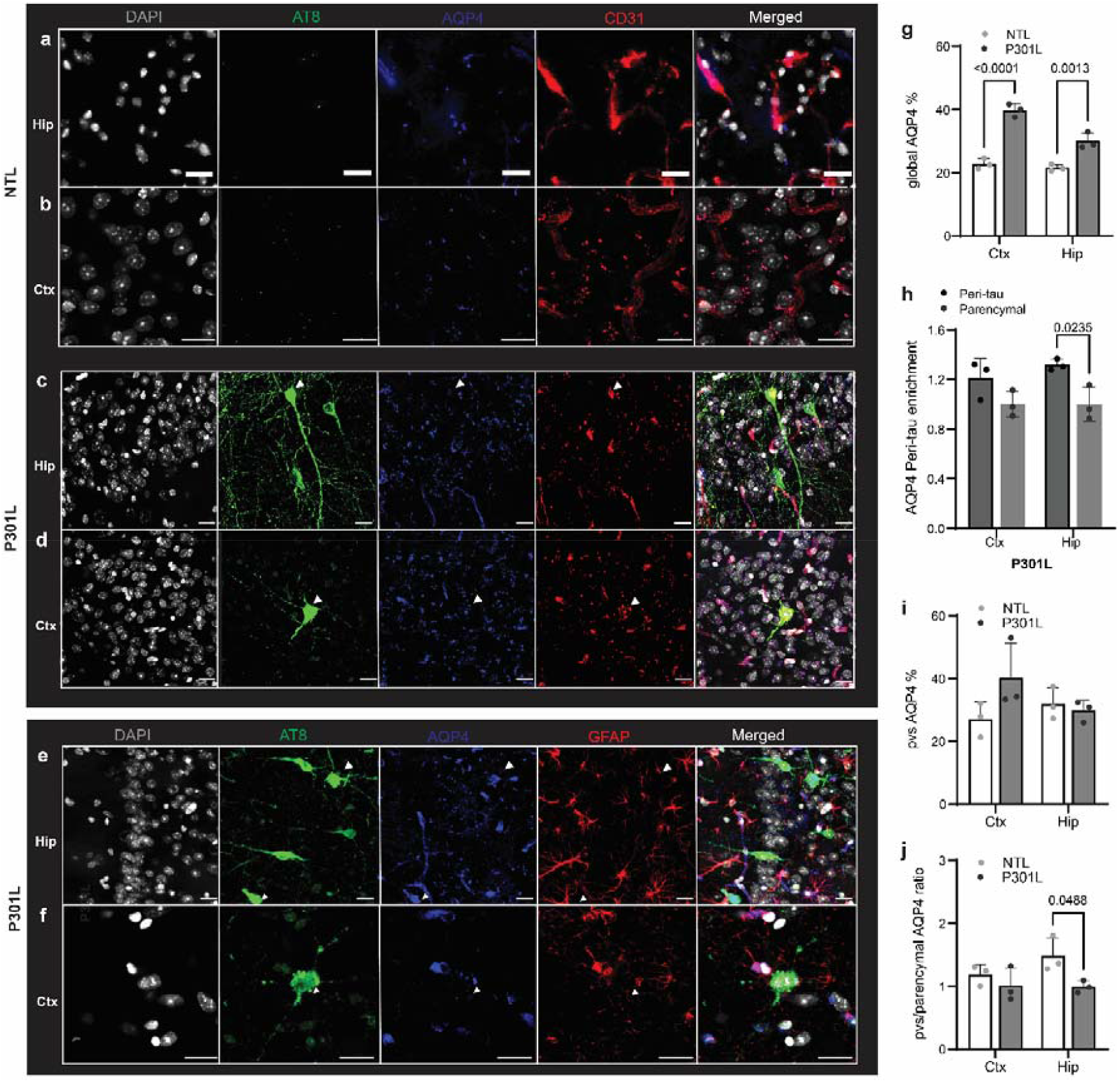
Increased AQP4 levels, depolarization and association with tau inclusions in P301L mouse brain. (**a-d**) Brain tissue sections of nontransgenic (NTL, n = 3) and P301L mice (n = 3) were stained for phospho-tau (AT-8 antibody, green), AQP4 (blue), and CD31 (red) in the hippocampus (Hip) and cortex (Ctx, forebrain). Nuclei were counterstained with DAPI (white). Scale bar = 20 μm. (**e-f**) Brain tissue sections of nontransgenic (NTL, n = 3) and P301L mice (n = 3) were stained for phospho-tau (AT-8 antibody, green), AQP4 (blue), and GFAP (red) in the Hip and Ctx. Nuclei were counterstained with DAPI (white). Scale bar = 20 μm. Overlapping of tau/AQP4/CD31 and tau/AQP4/GFAP was observed in mouse brain (white arrowhead in c-f). (**g**) Increased AQP4 signal in the Hip and Ctx of P301L compared to NTL. (**h**) Increased AQP4 signal peri-tau in the Hip of P301L mice. (**i**) No change in perivascular (pvs) AQP4 signal in the Hip and Ctx of P301L compared to NTL. (**j**) Reduced pvs/parenchymal AQP4 ratio in the Hip of P301L compared to NTL, indicating depolarization. Data are presented as the mean ± standard deviation.

We further analyzed the perivascular AQP4 immunofluorescence in mouse brain slices. No difference was observed in the perivascular absolute AQP4 level in the cortex (40.3 ± 11.1 vs 27.1 ± 5.4) or hippocampus (30.0 ± 3.4 vs 31.9 ± 5.1) of P301L mice (n = 3) compared to NTL mice (n = 3) (**Fig. 2i**). A reduced perivascular/parenchymal AQP4 ratio was observed in the hippocampus (1.0 ± 0.1 vs 1.5 ± 0.3, approx. 50 % reduction, p = 0.0488) of P301L (n = 3) compared to NTL mice (n = 3), indicating AQP4 depolarization (**Fig. 2j**). No difference was observed in the perivascular/parenchymal AQP4 ratio in the cortex (1.0 ± 0.3 vs 1.2 ± 0.2) of P301L mice (n = 3) compared to NTL mice (n = 3) (**Fig. 2j**).

### Gliosis in P301L mice

Next, we assessed astrogliosis in P301L mice by immunohistochemical staining using s100β-GFAP antibodies (**Fig. 3a-d**). The levels of s100β were increased in the cortex (2.1 ± 0.4 vs 0.6 ± 0.1, p < 0.0001) and hippocampus (2.1 ± 0.4 vs 0.5 ± 0.2, p < 0.0001) of P301L (n = 3) and NTL mice (n = 3) (**Fig. 3g**). The levels of GFAP were increased in the cortex (2.8 ± 0.5 vs 1.4 ± 0.5, p = 0.0118) and hippocampus (6.1 ± 0.3 vs 2.1 ± 0.6, p < 0.0001) of P301L (n = 4) and NTL mice (n = 4) (**Fig. 3h**). We assessed microgliosis in P301L mice by immunohistochemical staining using CD68-Iba1 antibodies (**Fig. 3e-f**). The levels of CD68 were increased in the cortex (2.3 ± 0.3 vs 1.2 ±0.5, p = 0.0113) but not in the hippocampus (2.4 ± 0.5 vs 1.6 ± 0.1) of P301L (n = 3) and NTL mice (n = 3) (**Fig. 3i**). The levels of Iba1 were increased in the cortex (0.7 ± 0.1 vs 0.2 ± 0.1, p = 0.0365) and hippocampus (1.0 ± 0.2 vs 0.4 ± 0.2, p = 0.0051) of P301L (n = 3) and NTL mice (n = 3) (**Fig. 3j**).

**Fig. 3.**
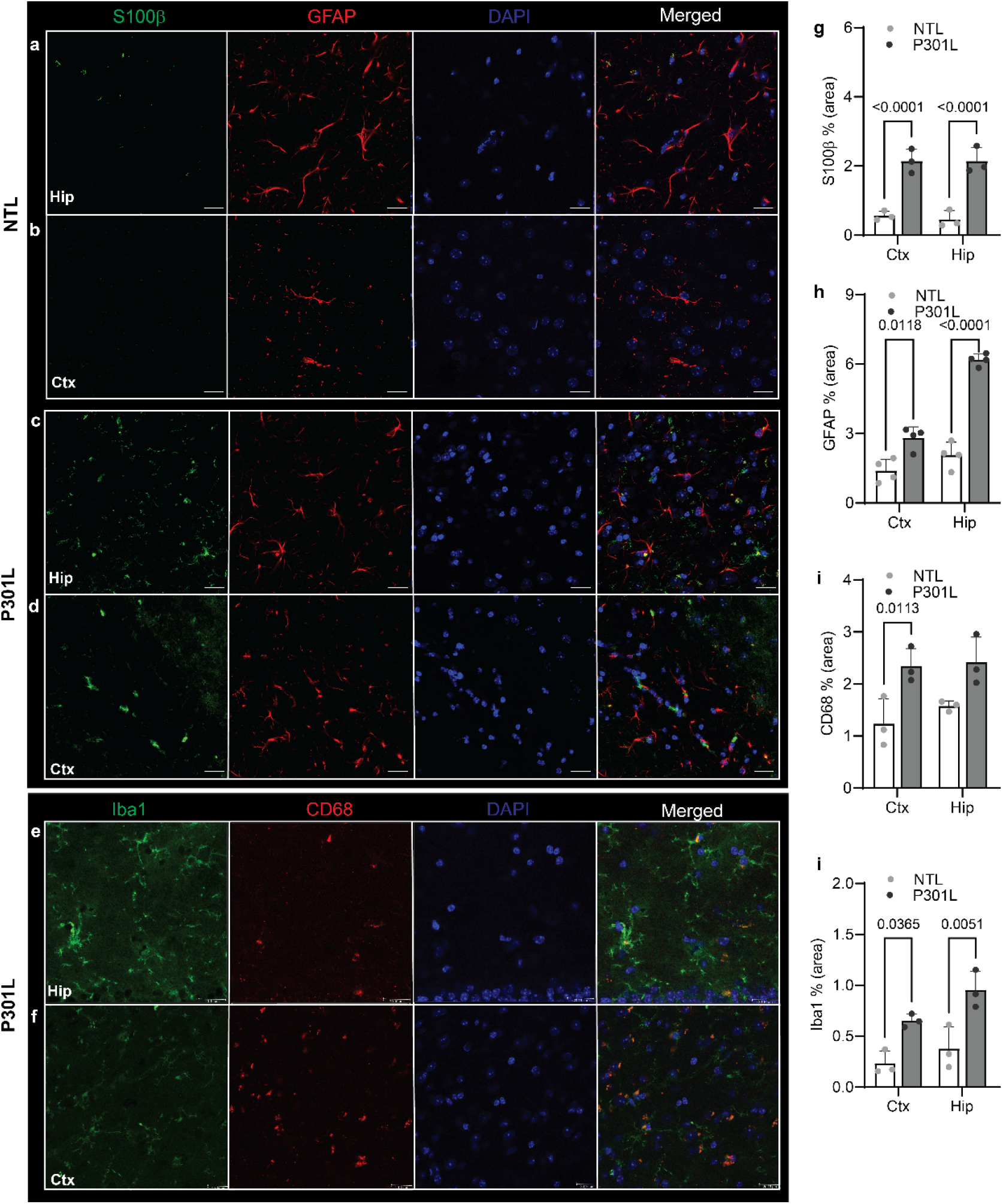
Increased astrogliosis and microgliosis in P301L mouse brain. (**a-d**) Brain tissue sections of nontransgenic (NTL, n = 3) and P301L mice (n = 3) were stained for S100β (green) and GFAP (red) in the hippocampus (Hip) and cortex (Ctx). Nuclei were counterstained with DAPI (blue). Scale bar = 20 μm. **(e-f**) Brain tissue sections of nontransgenic (NTL, n = 3) and P301L (n = 3) mice were stained for Iba1 (green) and CD68 (red) in the Hip) and Ctx. Nuclei were counterstained with DAPI (blue). Scale bar = 20 μm. **(g, h**) Increased S100β and GFAP signals in the Hip and Ctx of P301L compared to NTL. **(i, j**) Increased CD68 and Iba1 signals in the Ctx and increased Iba1 signals in the Hip of P301L compared to NTL.

### Increased number of CMBs in the brains of arcAβ mice

Next, we examined the neurovascular changes in the arcAβ mice. We observed hypointensities and paramagnetic lesions in the SWI images (validated by the hypointensities in the phase images) of arcAβ mice at 16 months of age, indicative of CMBs. The CMBs were distributed in both cortical and subcortical regions in the arcAβ mouse brains (**Fig. 4b**). Regional analysis showed a significantly greater number of CMBs in the brains of arcAβ (n = 8) than in the whole brains of NTL (n = 8) (49.9 ± 18.1 vs 0.25 ± 0.7, p = 0.0002) (**Fig. 4c**). Only 1 or 2 CMBs were found in the brains of NTL (**Figs. 4a**). Hematoxylin and eosin staining in the brain tissue slices of the arcAβ mice postmortem validated the presence of CMBs (**Fig. 4d**). Immunohistochemical staining using CD31 further showed an approx. 40 % reduction in the vessel density in the cortex (1.7 ± 0.3 vs 3.9 ± 0.1, p < 0.0001) and CA1 region of the hippocampus (1.6 ± 0.2 vs 3.7 ± 0.2, p < 0.0001) of arcAβ mice (n = 3) compared to NTL mice (n = 3) (**Figs. 4e, f, g**).

**Fig 4.**
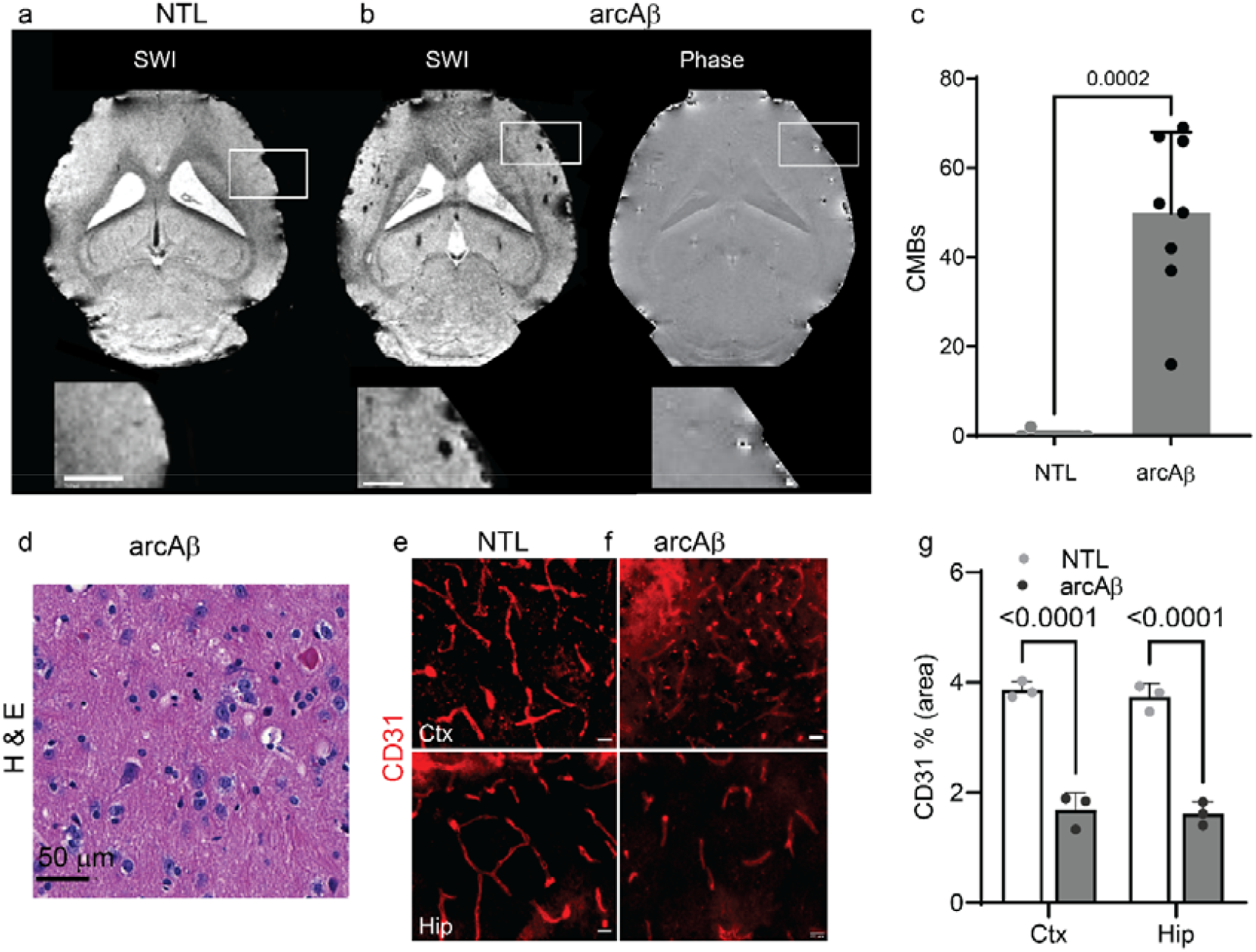
*In vivo* susceptibility weighted imaging (SWI) in the arcAβ mouse brain. (**a-c**) A representative *in vivo* SW image at 9.4 T revealed paramagnetic lesion hypointensities in the cortex and subcortical regions in arcAβ mice. Corresponding phase image of the arcAβ mouse brain showing hyperintensities/negative phase shifts indicating the presence of iron-rich cerebral microbleeds (CMBs). (**c**) Quantification of CMBs in arcAβ mice (n = 8) compared to nontransgenic littermates (NTL, n = 8). Mann–Whitney test, *** p < 0.001; (**d**) Representative hematoxylin and eosin (H & E) staining in the cortex of arcAβ mice showing the presence of CMBs. Scale bar = 50 μm. (**e-g**) CD31 staining showing reduced vessel density in the cortex and hippocampus of arcAβ mice compared to NTL mice. Scale bar = 20 μm. Data are presented as the mean ± standard deviation.

### Association of AQP4 with Aβ deposits and AQP4 polarization in the arcAβ mouse brain

Next, we performed triple staining using AQP4/CD31/6E10 (Aβ) and AQP4/GFAP/6E10 in the brain slices of arcAβ mice. The accumulation of Aβ deposits in both parenchymal plaques and CAA was detected (6E10 positive) across the cortex and hippocampus of arcAβ mice. No specific Aβ immunoreactivity was detected in the brains of NTL (**Figs. 5, SFigs. 2-4**). AQP4 staining was homogenous in the brain slices of NTL mice (**Figs. 5a, b**). The cortical (37.5 ± 7.1 vs 27.5 ± 5.3) and hippocampal (35.2 ± 2.0 vs 23.8 ± 4.9) levels of AQP4 immunofluorescence % (absolute value) were not significantly different between arcAβ (n = 3) and NTL mice (n = 3) (**Fig. 5g**). The AQP4 immunofluorescence intensity increased locally around the parenchymal plaques and CAA in the brains of arcAβ mice. The peri-Aβ/parenchymal ratio of AQP4 was approx. 1.6 ± 0.1 and 1.3 ± 0.02 in the cortex and hippocampus of arcAβ mice, respectively (**Fig. 5h**). Colocalization of Aβ as CAA with CD31-labelled endothelial cells and AQP4 was detected in the capillaries and arterioles of arcAβ mice (**Fig. 5c, d**). The patterns of AQP4-Aβ association were observed in the cortex and hippocampus of arcAβ mice, depending on the plaque status: 1) increase exclusively in the margin of the dense core plaques (**Figs. 5e, f**), and 2) distributed across the entire plaques (**Figs. 5c**).

**Fig. 5.**
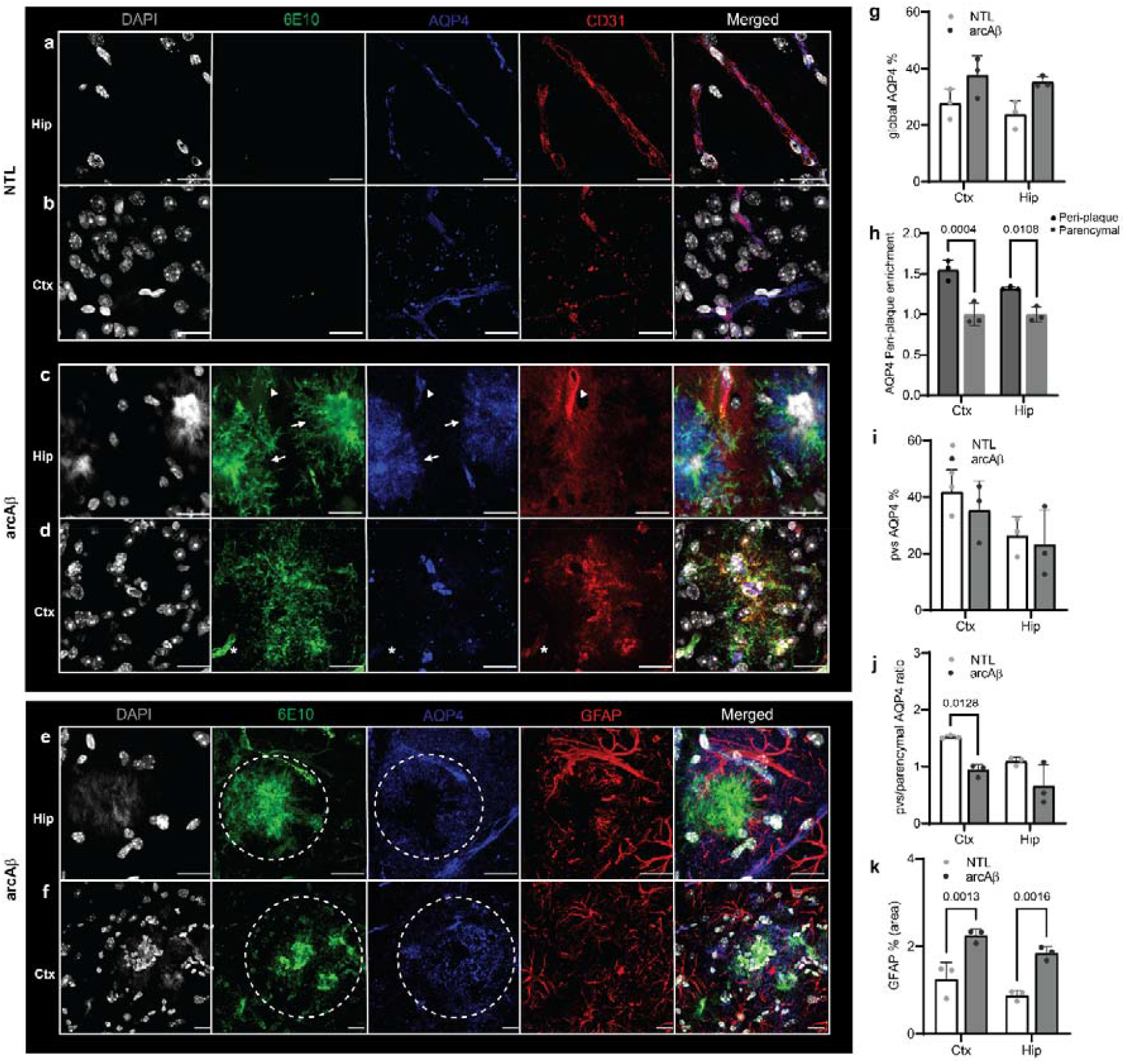
AQP4 depolarization and peri-plaque enrichment in arcAβ mice. (**a-d**) Brain tissue sections of nontransgenic (NTL, n = 3) and arcAβ mice (n = 3) were stained for Aβ (6E10 antibody, green), AQP4 (blue), and CD31 (red) in the hippocampus (Hip) and cortex (Ctx). Nuclei were counterstained with DAPI (white). Increased AQP4 immunoreactivity throughout the diffuse plaque (white arrow). Cerebral amyloid angiopathy (CAA) is indicated by a white arrowhead. Scale bar = 20 μm. CAA (overlap with AQP4/CD31) in the capillary indicated by white*. (**e, f**) Staining for Aβ (6E10, green), AQP4 (blue), and GFAP (red) in the Hip and Ctx. Nuclei were counterstained with DAPI (white). Dense plaque with AQP4 immunoreactivity at the margin (white circle). Scale bar = 20 μm. (**g**) No change in AQP4 signal in the Hip and Ctx of arcAβ compared to NTL. (**h**) Peri-Aβ plaque enrichment of AQP4 in the Ctx and Hip of arcAβ mice. (**i**) No change in perivascular (pvs) AQP4 signal in the cortex and hippocampus of arcAβ compared to NTL. (**j**) Reduced pvs/parenchymal AQP4 ratio in the Ctx of arcAβ compared to NTL, indicating depolarization. (**k**) Increased GFAP signal in the Ctx and Hip of arcAβ compared to NTL. Data are presented as the mean ± standard deviation.

We further analyzed the perivascular AQP4 immunofluorescence. The absolute perivascular AQP4 immunofluorescence level was not significantly different in the cortex (35.4 ± 10.4 vs 41.8 ± 7.8) and hippocampus (23.2 ± 12.3 vs 26.3 ± 6.8) between arcAβ (n = 3) and NTL mice (n = 3) (**Fig. 5i**). A 40 % reduction in the perivascular/parenchymal AQP4 ratio was observed in the cortex (0.9 ± 0.1 vs 1.5 ± 0.03, p = 0.0128) of arcAβ mice (n = 3) compared to NTL mice (n = 3), indicating AQP4 depolarization (**Fig. 5j**). No difference was observed in the perivascular/parenchymal AQP4 ratio in the hippocampus (0.7 ± 0.4 vs 1.1 ± 0.1) of arcAβ mice (n = 3) compared to NTL mice (n = 3). (**Fig. 5j**).

### Gliosis in arcAβ mice

The levels of GFAP were increased in the cortex (3.2 ± 0.1 vs 1.2 ± 0.4, p =0.0013) and hippocampus (1.8 ± 0.2 vs 0.9 ± 0.1, p = 0.0016) of arcAβ (n = 3) and NTL mice (n = 3) (**Fig. 5k**). Increased levels of GFAP were observed surrounding the Aβ plaques (**SFig. 3b**) and CAA (**SFig. 3c**) in the cortex and hippocampus of arcAβ mice compared to NTL mice. We further assessed the levels of microgliosis by immunohistological staining using Iba1-6E10 on brain slices from arcAβ mice. Activated microglia surrounding Aβ were observed in the cortex and hippocampus of arcAβ mice (**SFig. 4**).

## Discussion

Here, we demonstrated different patterns of neurovascular and AQP4 alterations in mouse models of tau and amyloidosis. Increased cortical and hippocampal AQP4 levels, hippocampal AQP4 depolarization, preserved rCBF and reduced cerebral vessel density were observed in P301L mice. In contrast, peri-plaque AQP4 enrichment, cortical AQP4 depolarization, reduced cerebral vessel density, and CMBs were detected in arcAβ mice.

We observed that Aβ and tau were differentially associated with AQP4. In pR5 P301L mice, AQP4 immunofluorescence was increased in the cortex and hippocampus (**Fig. 2**). AQP4 enrichment was detected peri-tau in the hippocampus of P301L mice (**Fig. 2**) and peri-Aβ deposits in both the cortex and hippocampus of arcAβ mice (**Fig. 5**). In addition, AQP4 depolarization appeared to be region-specific: in the hippocampus of P301L mice and in the cortex of arcAβ mice. Upregulated *AQP4* mRNA expression, immunofluorescence, and AQP4 depolarization were detected in the caudal cortex of rTg4510 mice [52]. The glymphatic system clears extracellular tau and protects against tau aggregation and neurodegeneration in PS19×AQP4 knockout mice [77]. In addition, altered neurons facing astrocytic AQP4 expression and glymphatic drainage of abnormal tau were found in IL33-deficient mice [78]. We found that dense parenchymal Aβ plaques showed increased AQP4 expression only at the margins, while AQP4 immunoreactivity was intense in CAA as well as in the interior of diffuse Aβ plaque (**Fig. 5**). The link between tau and AQP4 has been recently shown in two other tau mouse models. Our observed patterns of peri-plaque AQP4 enrichment in arcAβ mice were in agreement with previous reports in tg-ArcSwe mice [24, 79, 80], 5×FAD [74, 81, 82], and APP/PS1 [77, 83–87] mouse models. Behavioral and electrophysiological evidence indicated a neuroprotective role of AQP4 in 5×FAD mice [81].

We found impaired neurovasculature in both tau and Aβ mice. Reduced vessel density, preserved rCBF and colocalization of tau with CD31-stained endothelial cells were observed in pR5 P301L mice, and reduced vessel density with pronounced parenchymal Aβ/CAA was observed in arcAβ mice (**Figs. 1, 4**). Tau induces blood vessel abnormalities and angiogenesis-related gene expression in mice and in patients with AD [88, 89]. The alteration of rCBF in tau mouse models has been inconclusively reported. Park et al. reported that tau induced neurovascular uncoupling and reduced the activity-dependent rCBF in P301S mice (PS19) and P301L (rTg4510) mice compared to NTL [90]. Another study reported increased rCBF in P301L (rTg4510) mice [91]. Similar to previous observations in pR5 P301L mice [62], we observed preserved rCBF in pR5 P301L mice at 10-12 months of age, despite reduced cerebral vessel density and the presence of regional brain atrophy. The association of Aβ and rCBF has also been investigated extensively in other animal models of amyloidosis, such as APP/PS1 [92], and in patients with AD [93]. We have previously shown cerebrovascular alterations, including compromised rCBF, reduced cerebrovascular reactivity, and reduced vessel density in the cortex of arcAβ mice [54, 58, 59, 67, 94]. Aβ oligomers constrict capillaries in the brains of patients with AD by signaling to pericytes [95]. Our finding of SWI detection of CMBs in arcAβ mice is consistent with a previous MRI report in arcAβ mice [60], as well as in APP23 mice [58, 96]. CAA accumulation leads to CMBs and perfusion changes [97] and aggravates perivascular clearance impairment in a mouse model [98].

We observed increased levels of s100β and GFAP immunofluorescence, indicating astrogliosis, in the brains of both P301L and arcAβ mice compared to NTL mice (**Figs. 2, 3, 5, SFig. 3**), in line with previous reports [54, 99–102]. Reactive astrocytes have been shown to acquire neuroprotective and deleterious signatures in response to tau in PS19 mice and Aβ pathology in APP/PS1 mice [86]. Tau induced reactive astrocytes and loss of their neurosupportive functions in mouse models, including PS19, Thy-Tau22, hTau and rTgTauEC mice [52, 103–108]. Astrogliosis in turn could increase tau hyperphosphorylation and aggregation [109]. Astrocyte-neuronal network interplay is disrupted in APP/PS1 mice [110]. Removing senescent glial cells prevented tau-dependent pathology and cognitive decline in PS19 mice [111]. Early astrocytosis has been demonstrated by positron emission tomography tracers in both patients with AD and amyloidosis animal models [112–118]. Tau accumulates in hilar astrocytes of the dentate gyrus in postmortem brains from patients with AD [119–121] and induces neuronal dysfunction and memory deficits in a mouse model [122]. Hippocampal GFAP-positive astrocyte responses to Aβ and tau in postmortem brain tissue from patients with AD [123]. Moreover, senescent astrocytes were found near tangles in postmortem brain tissue from patients with AD and FTLD [124]. A glial profiling study demonstrated enrichment of astrocytic transcripts in tau-related FTLD degeneration [125]. In humans, tau immunotherapy results have been shown to be associated with glial responses in FTLD-tau [126].

There are several limitations of this study. First, we did not assess glymphatic function using *in vivo* imaging. Several MRI methods have been developed for glymphatic CSF and solute transport analysis [52, 127–129]. Second, we did not include the quantification of AQP4 mRNA expression in mouse brain tissues. Given that the expression of AQP4 was inhomogeneous and localized with Aβ and region-specific in both lines, mRNA analysis of the brain homogenate will likely average over the signal. Third, we did not assess the transcriptomic status of astrocytes in the mouse model. Further studies to investigate the transcriptomic signature associated with tau and Aβ will be informative.

In conclusion, we observed neurovascular alterations and divergent regional patterns of AQP4 increase and depolarization associated with phospho-tau and Aβ pathologies.

## Acknowledgement

The authors acknowledge Dr Mark Aurel Augath, Ms Diana Kindler, Dr Marie Roulaut at the Institute for Biomedical Engineering, ETH Zurich & University of Zurich; Ms Iva Mladenova, Ms Cinzia Maschio, Mr Daniel Schuppli at the Institute for Regenerative Medicine, University of Zurich for technical assistance.

## Author contribution statement

RN designed the study; VK performed the staining and microscopy; LB and RN performed MRI; VK, JK and RN analysed and interpreted the data. VK, JK and RN wrote the first draft. All authors contributed to the manuscript.

## Ethics declaration

All experiments were performed in accordance with the Swiss Federal Act on Animal Protection and were approved by the Cantonal Veterinary Office Zurich (permit number: ZH082/18, ZH162/20).

