## Supplementary materials for "Aquaporin 4 is differentially increased and depolarized in association with tau and amyloid-beta"

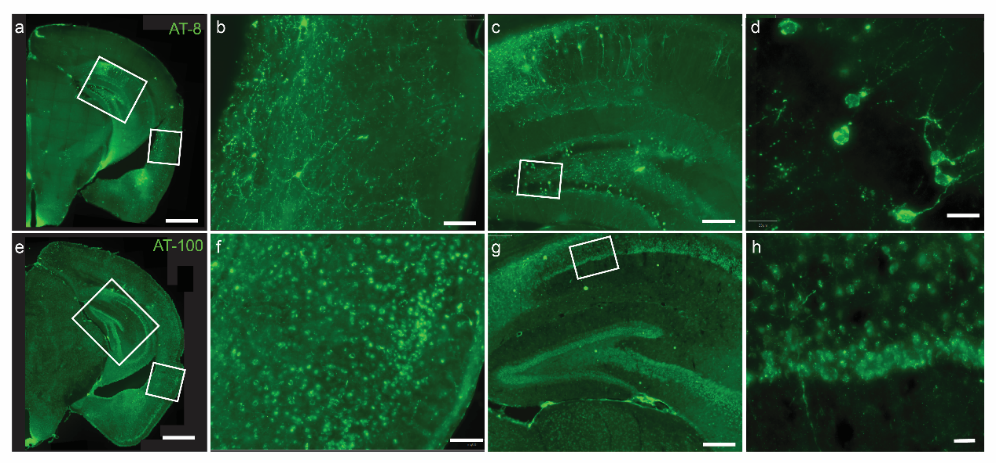

**Supplementary Fig. 1 Tau inclusions in the forebrain and hippocampus of P301L mice** (**a-h**) Immunofluorescence staining using anti-phospho-tau antibodies AT-8 (a-d) and AT-100 (e-h) on brain slices from P301L mice. Tau inclusions were detected mainly in the forebrain (b. f) and hippocampus (c, d, zoom-in g, h). Hippocampal atrophy was observed in P301L mouse brain. Scale bar = 1 mm (a, e), 100 μm (b, f), 200 μm (c, g), 20 μm (d, h).

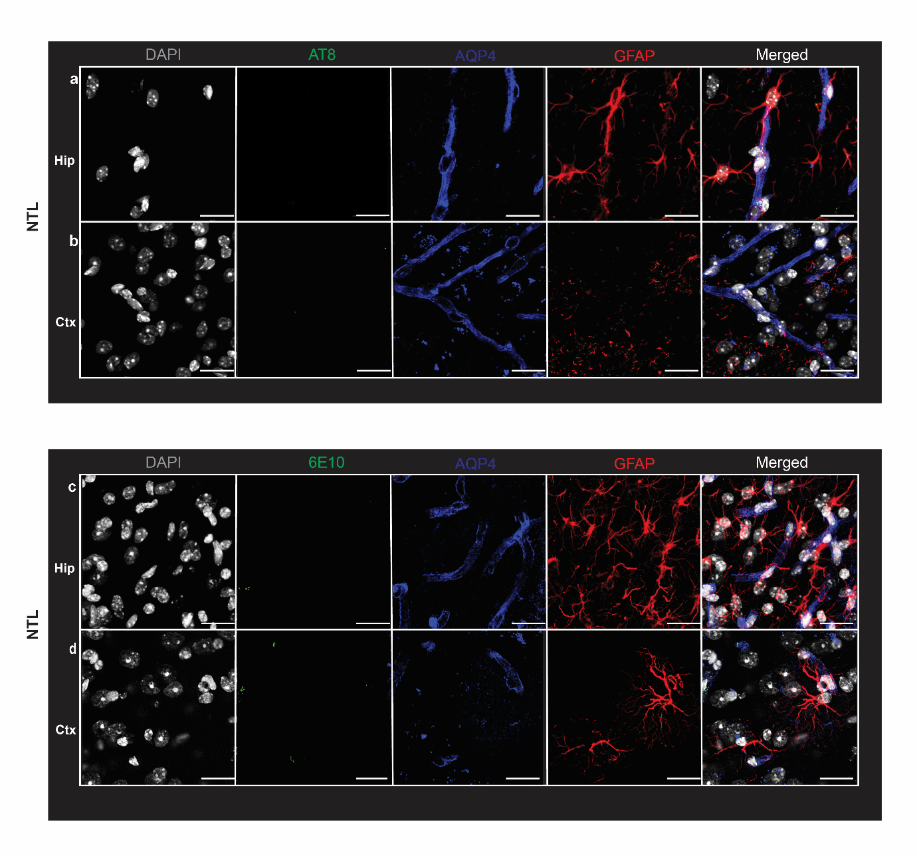

**Supplementary Fig. 2 (a-b**) Brain tissue sections of nontransgenic (NTL) of P301L mice were stained for phospho-tau (AT-8 antibody, green), AQP4 (blue), and GFAP (red) in the hippocampus (Hip) and cortex (Ctx). Nuclei were counterstained with DAPI (white). Scale bar = 20 μm. **(c-d**) Brain tissue sections of nontransgenic (NTL) of arcAβ mice were stained for amyloid-beta (6E10 antibody,green), AQP4 (blue), and GFAP (red) in the hippocampus (Hip) and cortex (Ctx). Nuclei were counterstained with DAPI (white). Scale bar = 20 μm.

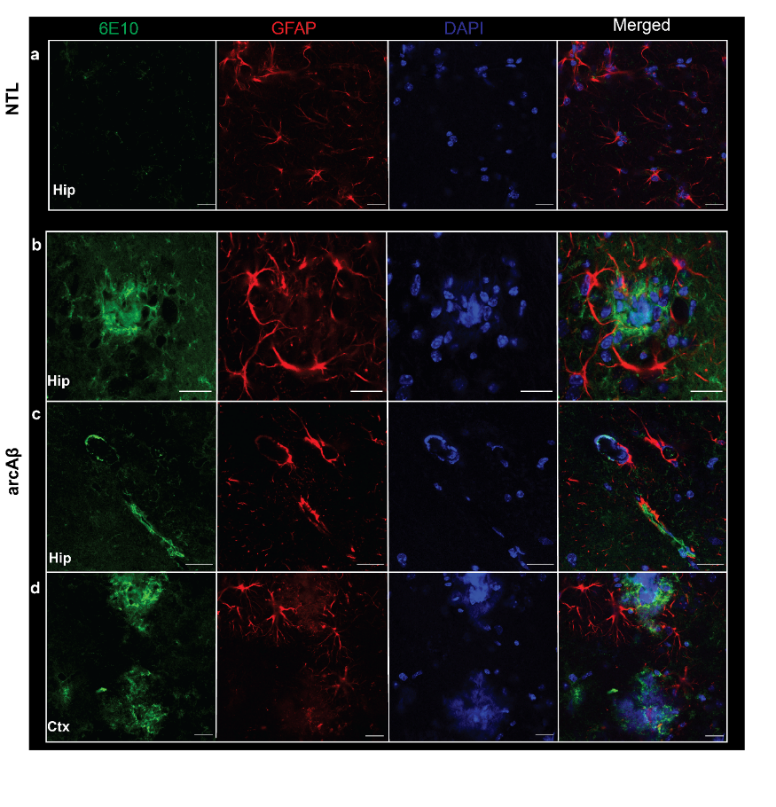

**Supplementary Fig. 3 Increased** **GFAP in the cortex and hippocampus of arcAβ mice. (a-d**) Brain tissue sections of nontransgenic (NTL, n = 3) and arcAβ mice (n = 3) were stained for Aβ (6E10 antibody, green), and GFAP (red) in the hippocampus (Hip) and cortex (Ctx). (Nuclei were counterstained with DAPI (blue). Parenchymal plaque (b, d) and cerebral amyloid angiopathy (c) were observed in arcAβ mice. Scale bar = 20 μm. Data are presented as the mean ± standard deviation.

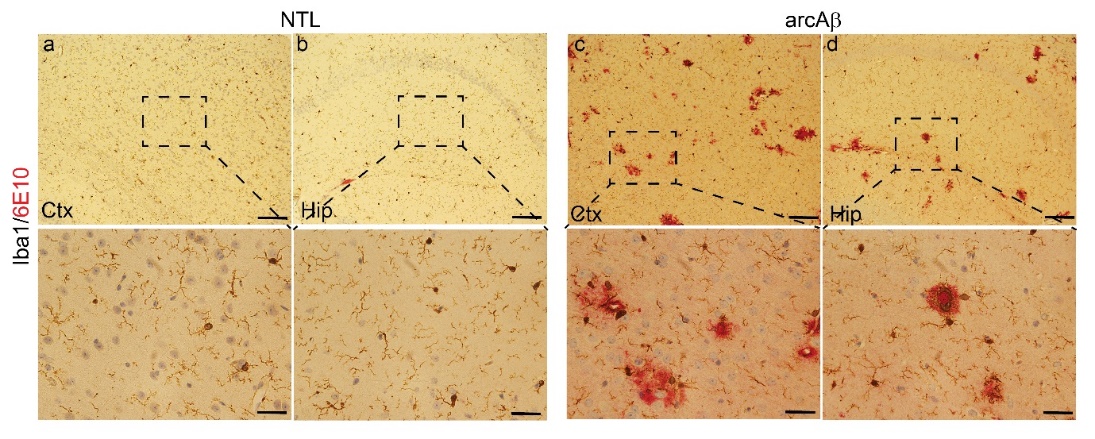

**Supplementary Fig. 4** **Immunohistochemical analysis of Aβ load and microglia activation.** (A) Brain tissue sections of NTLs and arcAβ mice were stained for amyloid (6E10, red), microglia (Iba1, black) immunoreactivity in the cortex (Ctx) and hippocampus (Hip). Scale bar = 100 μm (upper panel), 25 μm (lower panel).

**Supplementary table 1 Summary of antibodies and chemicals used in staining**

|  | **Catalog no** | **Dilution** | **Supplier** |
| --- | --- | --- | --- |
| AT-8 | MN1020 | 1:1000 | Invitrogen |
| AT-100 | MN1060 | 1:1000 | Invitrogen |
| 6E10 | 803001 | 1:1000 | Biolegend |
| AQP4 | HPA014784 | 1:1000 | Atlas |
| CD31 | 550274 | 1:1000 | BD |
| GFAP | PAA056MU01 | 1:1000 | Cloud Clone |
| S100β | 710363 | 1:1000 | Invitrogen |
| Iba1 | PA527436 | 1:1000 | Invitrogen |
| CD68 | Ab213363 | 1:1000 | Abcam |
| Alexa-488 | A32723 | 1:200 | Invitrogen |
| Alexa-647 | A32795 | 1:200 | Invitrogen |
| VECTASHIELD fluorescent Antifade mounting media | H-1000-10 |  | Vector Laboratories |
| Normal goat Serum | JAC005-000-121 |  | Chemie Brunschwig AG |
| Normal donkey Serum | R37624 |  | Invitrogen |
| DAPI | D1306 | 1:1000 | Sigma-Aldrich |
